# Visualization of early RNA replication kinetics of SARS-CoV-2 by using single molecule RNA-FISH

**DOI:** 10.1101/2022.12.10.517707

**Authors:** Rajiv Pathak, Carolina Eliscovich, Ignacio Mena, Updesh Dixit, Adolfo García-Sastre, Robert H Singer, Ganjam V. Kalpana

**Affiliations:** Departments of Genetics and Microbiology and Immunology, Albert Einstein College of Medicine, Bronx, New York, USA; Departments of Medicine (Hepatology), and Developmental and Molecular Biology, Albert Einstein College of Medicine, Bronx, New York, USA; Department of Microbiology, Icahn School of Medicine at Mount Sinai, New York, USA; Global Health and Emerging Pathogens Institute, Icahn School of Medicine at Mount Sinai, New York, USA; Department of Medicine, Division of Infectious Diseases, Icahn School of Medicine at Mount Sinai, New York, USA; The Tisch Cancer Institute, Icahn School of Medicine at Mount Sinai, New York, USA; Department of Pathology, Molecular and Cell-Based Medicine, Icahn School of Medicine at Mount Sinai, New York, USA; Departments of Cell Biology and Neuroscience, Albert Einstein College of Medicine, Bronx, New York, USA

**Author notes:** Corresponding author: Ganjam V. Kalpana, Ph.D., Mark Trauner Faculty Scholar in Neuro-Oncology, Professor, Department of Genetics, Professor, Department of Microbiology, and Immunology, Albert Einstein College of Medicine, 1300, Morris Park Ave., Ullman 821, Bronx, New York, 10461.

## Abstract

SARS-CoV-2 infection has caused a major global burden. Despite intensive research, the mechanism and dynamics of early viral replication are not completely understood including the kinetics of formation of plus stranded genomic and subgenomic RNAs (gRNA and sgRNA) starting from the RNA from the first virus that enters the cell. We employed single-molecule RNA-fluorescence *in situ* hybridization (smRNA-FISH) to simultaneously detect viral gRNA and sgRNA in infected cells and carried out a time course analysis to determine the kinetics of their replication. We visualized the single molecules of gRNA within the cytoplasm of infected cells 30 minutes post-infection and detected the co-expression of gRNA and sgRNA within two hours post-infection. Furthermore, we observed the formation of a replication organelle (RO) from a single RNA, which led to the formation of multiple ROs within the same cells. Single molecule analysis indicated that while gRNA resided in the center of these ROs, the sgRNAs were found to radiate and migrate out of these structures. Our results also indicated that after the initial delay, there was a rapid but asynchronous replication, and the gRNA and sgRNAs dispersed throughout the cell within 4-5 hours post-infection forming multiple ROs that filled the entire cytoplasm. These results provide insight into the kinetics of early post-entry events of SARS-CoV-2 and the formation of RO, which will help to understand the molecular events associated with viral infection and facilitate the identification of new therapeutic targets that can curb the virus at a very early stage of replication to combat COVID-19.

**Author Summary:** SARS-CoV-2 infection continues to be a global burden. Soon after the entry, SARS-CoV-2 replicates by an elaborate process, producing genomic and subgenomic RNAs (gRNA and sgRNAs) within specialized structures called replication organelles (RO). Many questions including the timing of multiplication of gRNA and sgRNA, the generation, subcellular localization, and function of the ROs, and the mechanism of vRNA synthesis within ROs is not completely understood. Here, we have developed probes and methods to simultaneously detect the viral gRNA and a sgRNA at single cell single molecule resolution and have employed a method to scan thousands of cells to visualize the early kinetics of gRNA and sgRNA synthesis soon after the viral entry into the cell. Our results reveal that the replication is asynchronous and ROs are rapidly formed from a single RNA that enters the cell within 2 hours, which multiply to fill the entire cell cytoplasm within ~4 hours after infection. Furthermore, our studies provide a first glimpse of the gRNA and sgRNA synthesis within ROs at single molecule resolution. Our studies may facilitate the development of drugs that inhibit the virus at the earliest possible stages of replication to minimize the pathogenic impact of viral infection.

## Introduction

Coronavirus disease 2019 (COVID-19) is a viral respiratory disease that emerged at the end of 2019 in Wuhan, China, and rapidly extended its devastating effects worldwide. COVID-19 is caused by SARS-CoV-2, a large enveloped, positive-strand RNA virus with a genome approximately 30 kb in length, belonging to the genus *Betacoronavirus* of the family Coronaviridae [1]. As of 16^th^ November 2022, SARS-CoV-2 continues to spread worldwide, with over 633.60 million total confirmed cases and 6.59 million deaths worldwide [2].

SARS-CoV-2 primarily targets the respiratory tract, and the infection proceeds with the binding of viral spike protein to the human receptor angiotensin-converting enzyme (ACE2), where spike cleavage by the transmembrane protease serine 2 (TMPRSS2) on the surface of epithelial cells triggers the fusion of the viral and host cell membranes [3, 4]. In addition to causing acute respiratory distress syndrome, SARS-CoV-2 has other pulmonary and extrapulmonary manifestations in the gastrointestinal tract, hepatobiliary system, cardiovascular, neurological, and renal systems, which often lead to multiorgan failure and shock in severe cases [5, 6]. Furthermore, some survivors experience Long COVID or Post-Acute Sequelae of SARS-CoV-2 (PASC), with cardiovascular, neurological, and pulmonary manifestations [7, 8]. The exact pathogenesis of acute and chronic disease in extrapulmonary organs in COVID-19 is unknown. It has been suggested that indirect mechanisms such as co-morbidities and/or other pathophysiological conditions may play a role [9, 10]. Understanding the details of mechanism of replication and cell tropism of SARS-CoV-2 may provide insights into the pathogenesis and tissue tropism of this virus.

*Coronaviridea* includes enveloped, positive sense, single-stranded RNA viruses. These viruses employ an elaborate mechanism for replicating their genome and for transcribing the coding sequences. The replication cycle of SARS-CoV-2, like other coronaviruses, begins with the entry of the virus into the cell and release of viral RNA into the cytoplasm [11]. Within the cytoplasm, the genomic positive-strand viral RNA (gRNA) acts as a template for translation of viral enzymes, including the viral RNA-dependent RNA polymerase (RdRP), and for the generation of negative-strand viral RNA replication intermediates. The newly synthesized negative strand RNA, in turn, acts as a template for generation of additional +strand gRNAs and a series of shorter sub-genomic RNAs (sgRNAs) by discontinuous transcription [11]. The structural and accessory proteins are synthesized from the shorter sgRNAs [1]. During its replication, the virus modifies the intracellular host endoplasmic reticulum membrane to generate the replication organelles (ROs), which are the powerhouses consisting of double-membrane vesicles (DMVs) that enclose the viral RNAs [12, 13]. Many questions remain unanswered about the early replication events of SARS-CoV-2 such as: i) the time it takes for the viral RNA to complete translation and start the replication process after entry of the viral RNA in to the cytoplasm; ii) the generation, subcellular localization, and function of the ROs; iii) timing of formation of gRNA and sgRNAs; and iv) the mechanism of vRNA synthesis within ROs [11]. Addressing these questions is not only important for obtaining insight about the mechanism of SARS-CoV-2 replication but also for identification of unexplored drug targets that can be used to curb the viral replication at a very early stage of the infection cycle.

Studies to understand the SARS-CoV-2 replication generally have utilized a high multiplicity of infection (m.o.i.) of the virus and have employed time points where viral replication is at a mid-point, where it can be easily monitored at ~4-5 hours post infection (p.i.) [14, 15]. These studies miss the replication events that occur very early (before 4-5 hours p.i.) after a single virus infectious particle enters the cell. Understanding of the early events of viral replication requires highly sensitive and specific methods to detect SARS-CoV-2 RNA at a single molecule level. In addition, when the viral replication has reached a mid-point by the time of analysis, it is generally not clear at what point the virus begins to replicate its +strand genome into multiple copies after the entry and at what point it begins to transcribe its sgRNAs.

To detect and visualize SARS-CoV-2 gRNA and sgRNAs with high specificity and sensitivity at very early time points, we employed a combination of single molecule RNA-Fluorescence in situ hybridization (smRNA-FISH). To facilitate the study of replication kinetics of a single SARS-CoV-2 RNA after it enters a cell we infected the cells with low m.o.i. of virus. Furthermore, we employed <u>H</u>igh-<u>S</u>peed <u>H</u>igh-<u>R</u>esolution <u>S</u>canning fluorescence microscopy (HSHRS-FM) to scan and visualize the replication of SARS-CoV-2 in a large number of cells by smRNA-FISH analysis. In HSHRS-FM method the instrument scans the entire slide creating a single high resolution digital image by tiling and stitching many high-magnification fields of view together, thus capturing the images of a large number of cells present on the slide. We designed probes to simultaneously detect positive strands of the gRNA and a sgRNA and carried out a time course analysis to visualize the cells at time points starting from 0.5 hours to 24 hours p.i. Our analyses have led to the detection of the SARS-CoV-2 gRNA within the infected cells as early as 30 minutes p.i. and allowed us to observe the earliest formation of individual RO from a single gRNA that entered the cell. We found that these RO contain both gRNA and sgRNAs and that the sgRNAs migrate out of these structures at early time points. Furthermore, we observed that multiple daughter ROs are formed from gRNAs that migrate out of the single parental RO arising from the virus that entered the cells to gives rise to newer ROs. These ROs ultimately fill the entire cell as the time progresses. We observed cells containing gRNA and sgRNA and ROs at various stages of their replication at any given time point, indicating that the replication is asynchronous. Our studies indicate that combining smRNA-FISH with HSHRS-FM will allow the sensitive detection of SARS-CoV-2 RNAs in a large number of infected cells to address emerging questions related to the mechanism of early replication events of SARS-CoV-2.

## Results

### Design and optimization of smRNA-FISH probes to study the early replication events of SARS-CoV-2 at single cell and single molecule resolution

To understand the kinetics and spatio-temporal aspects of the SARS-CoV-2 replication, we adopted smRNA-FISH method to facilitate the sensitive detection of gRNA and sgRNA-S at the single cell, single molecule level. Forty fluorescent labeled antisense oligonucleotides were synthesized based on the sequence of Spike gene, which would detect positive strands of gRNA and sgRNA-S (Fig 1A; see Methods).

**Fig 1.**
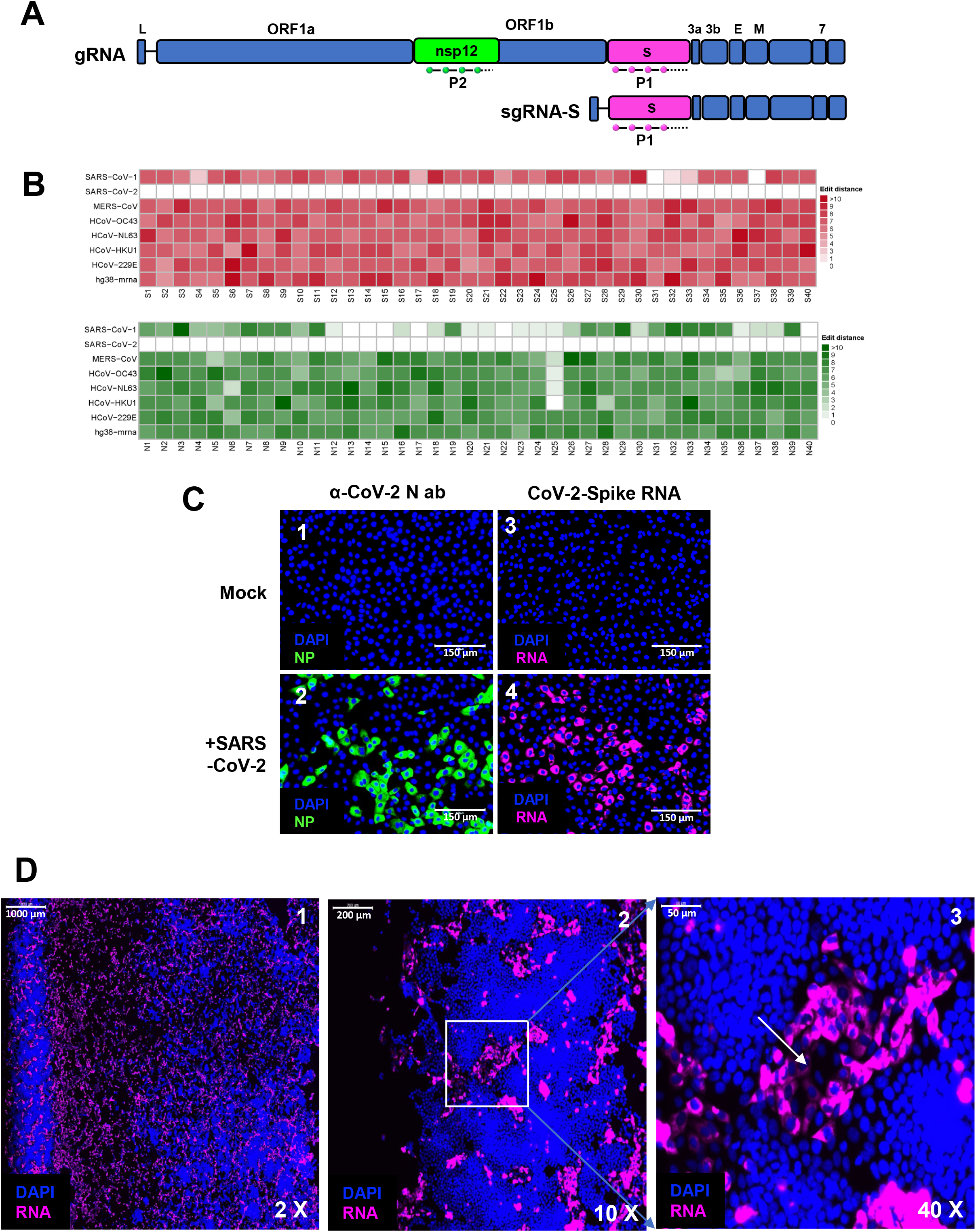
Design and optimization of SARS-CoV-2 specific smRNA-FISH probes to detect SARS-CoV-2 infection. **(A)** Schematic representation of SARS-CoV-2 gRNA and sgRNA-S, indicating the position of the smRNA-FISH probes, P1 (in magenta) and P2 (in green) probes that hybridize to Spike and nsp12 ORF sequences, respectively. **(B)** Heatmap representing the probe sequence alignment against various coronavirus genomes (including SARS-CoV-1, SARS-CoV-2, MERS-CoV, HCoV-OC43, HCoV-NL63, HCoV-229E) and human transcriptomes (hg38-mrna). Each column represents individual 22 nt spike gene probe sequences from Spike gene (S1-S40, upper panel, represented in the shades of red) and from nsp12 gene (N1-N40, lower panel, represented in shades of green). The minimum edit distance represents mismatch score between the SARS-CoV-2 sequence and the other genome sequences, where ‘0’ indicates a perfect match (in white) and >10 represents the most mismatch (in dark red or green). **(C)** Specificity of SARS-CoV-2 spike RNA Probe P1. Vero E6 cells mock- or SARS-CoV-2-infected, at 12 hours p.i. (m.o.i.: 0.5 PFU/cell), were probed with α-NP antibody (green, panels 1 and 2) as positive control for infection and spike RNA probe P1 (magenta, panels 3 and 4), respectively, to test the specificity of the probe. Blue color indicates nuclei upon DAPI staining. Scale bar, 150 μm. **(D)** Photomicrographs of scanned images of infected cells to determine the extent of SARS-CoV-2 infection and plaque formation. Vero E6 cells infected with SARS-CoV-2 virus at 0.5 m.o.i., were probed with spike RNA probe P1at 24 hours p.i. The entire slide was scanned at high speed and high resolution. The panels 1-3 represent images at 2x, 10x and 40x magnifications, respectively. The white arrow in the panel 3 represents the dead center of a plaque. Blue color represents DAPI staining of nuclei and Magenta, SARS-CoV-2 RNA. Scale bars, 1000 μm, 200 μm and 50 μm for 2x, 10x, and 40x images, respectively.

The specificity of oligonucleotides to SARS-CoV-2 sequences were analyzed by determining the edit distance between the oligonucleotides to the sequences of related human coronaviruses including SARS-CoV-1, MERS-CoV, HCoV-OC43, HCoV-NL63, HCoV-HKU1, HCoV-229E and to the human transcriptome, hg38-mRNA (Fig 1B, see Methods). Our analysis indicated that the oligonucleotide probes are specific to SARS-CoV-2 but not to the other coronaviruses or human sequences.

To determine the specificity of the probes to detect replicating SARS-CoV-2 RNA in cells, Vero E6 cells infected with <0.5 m.o.i. of infectious virus (SARS-CoV-2 USA-WA1/2020, GenBank MN985325.1) was subjected to smRNA-FISH using sgRNA-S probe, termed P1 (Fig 1C, panels 3 and 4). As a positive control, cells were independently analyzed by immunofluorescence (IF) using α-N antibodies to detect viral N protein expression (Fig 1C, panels 1 and 2). RNA-FISH probes and α-N antibodies detected SARS-CoV-2 RNA and N protein, respectively, in infected cells but not in the uninfected cells, establishing the specificity of the RNA-FISH probes (Fig 1C). As a negative control, we also infected Vero E6 cells with a VSV virus containing the sequence of codon optimized Spike gene open reading frame (VSV-Spike) [16]. We found that P1 probe did not give any positive signal with cells infected with VSV-Spike, indicating that the probes specifically detected wild type SARS-CoV-2 Spike gene sequences (S1 Fig).

Coronavirus is a lytic virus, and its spreading infection leads to the formation of plaques that harbor dead cells in the center and newly infected cells in the periphery [17]. To visualize the extent of viral replication, the infected Vero E6 cells were subjected to smRNA-FISH using the P1 probe 24 hour p.i., and the entire slide containing the infected cells was scanned at 20x magnification using HSHRS-FM. We found that at higher magnification, several small plaques characterized by the presence of groups of brightly fluorescent cells positive for P1 probe surrounding a patch of dead area were visible (Fig 1D). These studies indicated that the P1 probe selectively detects viral RNA in the infected live cells.

### Analysis of the kinetics of SARS-CoV-2 RNA replication

Upon viral entry into the cell, the first step is the synthesis of virally encoded critical enzymes including polymerases required for replication using the viral gRNA as a template. While most of the studies have investigated SARS-CoV-2 replication starting from ~4 hours p.i., where the replication can be visualized, one of the studies using smRNA-FISH analysis detected RNA within infected cells at 2 hours p.i [14]. To determine if we can detect the viral RNA at earlier times., we carried out a time course analysis starting from 0.5 hour p.i. To avoid multiple virus particle entry into the same cell, ~0.5 m.o.i. of the SARS-CoV-2 virus was used to infect cells (Fig 2A). The entire slide (>100,000 cells) with infected cells were imaged using HSHRS-FM at 20x to determine the percentage of cells infected. We observed that at ~6 hours p.i. ~7% of cells expressed brightly fluorescent RNA at this magnification (Fig 2A; panels 7-8). At 12 hours p.i. ~23% of the cells were infected and by 24 hours p.i. >50% of the cells were infected, reaching the numbers close to the estimated m.o.i. These results suggested that there is a delay in viral replication at early time points as only a small % of cells are scored as infected but at a later time point, viral replication can be easily visualized in 50% of the infected cells when infected with 0.5 m.o.i. of the virus (Fig 2A; panels 11-12).

**Fig 2.**
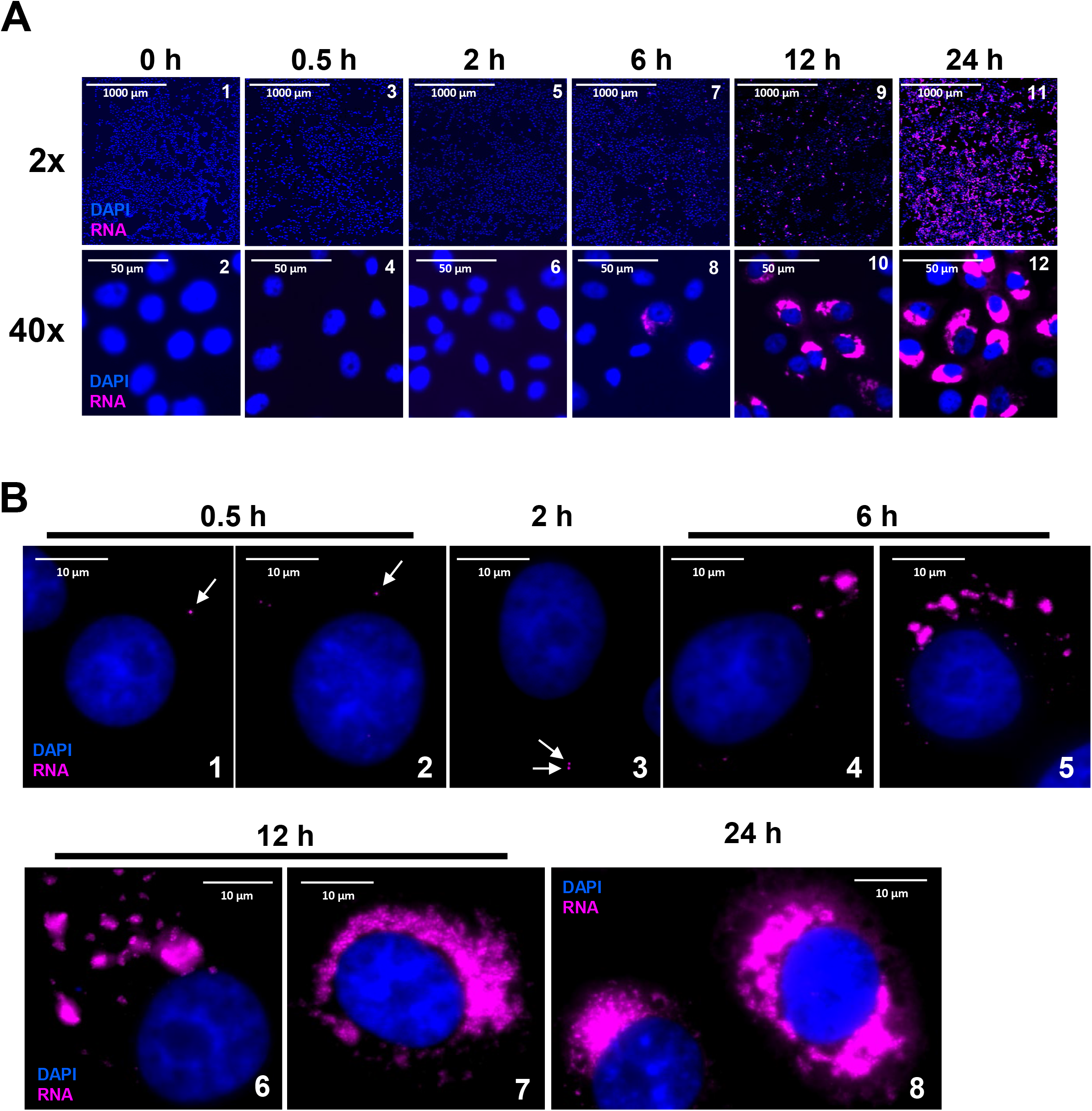
Time-course analysis of SARS-CoV-2 replication using smRNA-FISH. Cells were infected and hybridized with Quasar 570-labeled P1 probes (magenta) against spike RNA at 0, 0.5, 2, 6, 12 and 24 hours p.i. **(A)** Photomicrographs representing the images acquired using high resolution-high speed scanning microscopy are represented at 2x (upper panels) and 40x (lower panels) magnifications, respectively. Scale bars, 1000 μm and 50 μm for 2x and 40x images, respectively. **(B)** Photomicrographs representing images of infected cells at single cell single molecule resolution, probed with P1 at 0.5, 2, 6, 12 and 24 hours p.i., acquired using a wide-field Olympus BX-63 Microscope. White arrows point to single molecules of SARS-CoV-2 RNA seen at early time points, 0.5 and 2 hours p.i. DAPI (nucleus, blue). Scale bar, 10 μm.

To visualize early events of infection, the above samples were imaged using a fluorescence microscope fitted with a wide angled lens at 60x magnification and 1.4 numerical aperture (N.A.) oil objective that allows the detection of single viral RNA molecules within the infected cells. Since the Stellaris FISH probe comprises of 40 tandem 22-mer oligonucleotides corresponding to contiguous sequences on an mRNA, each with a fluorescent label, these probes collectively bind along the same target transcript to produce a punctate signal that appears as bright, diffraction-limited, computationally identifiable fluorescent spots [18–20]. The large number of probes in a Stellaris FISH assay ensures a high level of sensitivity and specificity and minimizes false negatives and background signals. We found that cells at 0.5 and 2 hours p.i. exhibited single molecules of viral RNA, which were not detected in uninfected samples (Fig 2B, panels 1-3, white arrows). Furthermore, we could not observe infected cells at 0.5 and 2 hours p.i. when using low magnification lens (Fig 2A). Analysis of Z-stacks of images revealed that the RNA signals in these images were detected only in the stacks that corresponded to interior of the cell and not the periphery, suggesting that these spots represent the earliest genomic RNA that entered the cell (S2A Fig). Furthermore, we found that some cells might have SARS-CoV-2 RNA within the nucleus at early time points (S2B Fig). At 6 hours p.i., spots that may represent single molecules as well as larger patches that represent groups of multiple RNA molecules were visible (Fig 2B, panel 4-5). Most of the FISH signals within the cells corresponded to patches of RNA rather than single RNA molecules at 6 hours p.i., making it difficult to quantitate the number of individual RNA molecules at these later time points (Fig 2B; panels 4 and 5). By 12 hours p.i., the size and number of these patches increased, and in most of the cells, the entire cytoplasm was filled with viral RNA (Fig 2B panels 6-7).

### Appearance of replication organelles (RO) during early stages of SARS-CoV-2 viral replication

Since spike RNA probe P1 detects both gRNA and sgRNA-S, we tested the ability of a second probe (P2) derived from *nsp12* gene sequences within ORF1b to detect the gRNA alone. The specificity of P1 and P2 probes were tested by carrying out smRNA-FISH analysis using either P1 or P2 probe sets or the two probe sets together and by analyzing the images obtained in both TritC (P1 probe) and Cy5 (P2 probe) channels (Fig 3A). We found that the probes were specific and there was no bleeding from one channel to the other (Fig 3A; panels 1-6). When the two probes were used together, signals from both probes were detected in the same set of infected cells (Fig 3A; panels 7-9). Neither one of the probes detected any signal in mock infected cells, confirming probe specificity (Fig 3A; panels 10-12).

**Fig 3.**
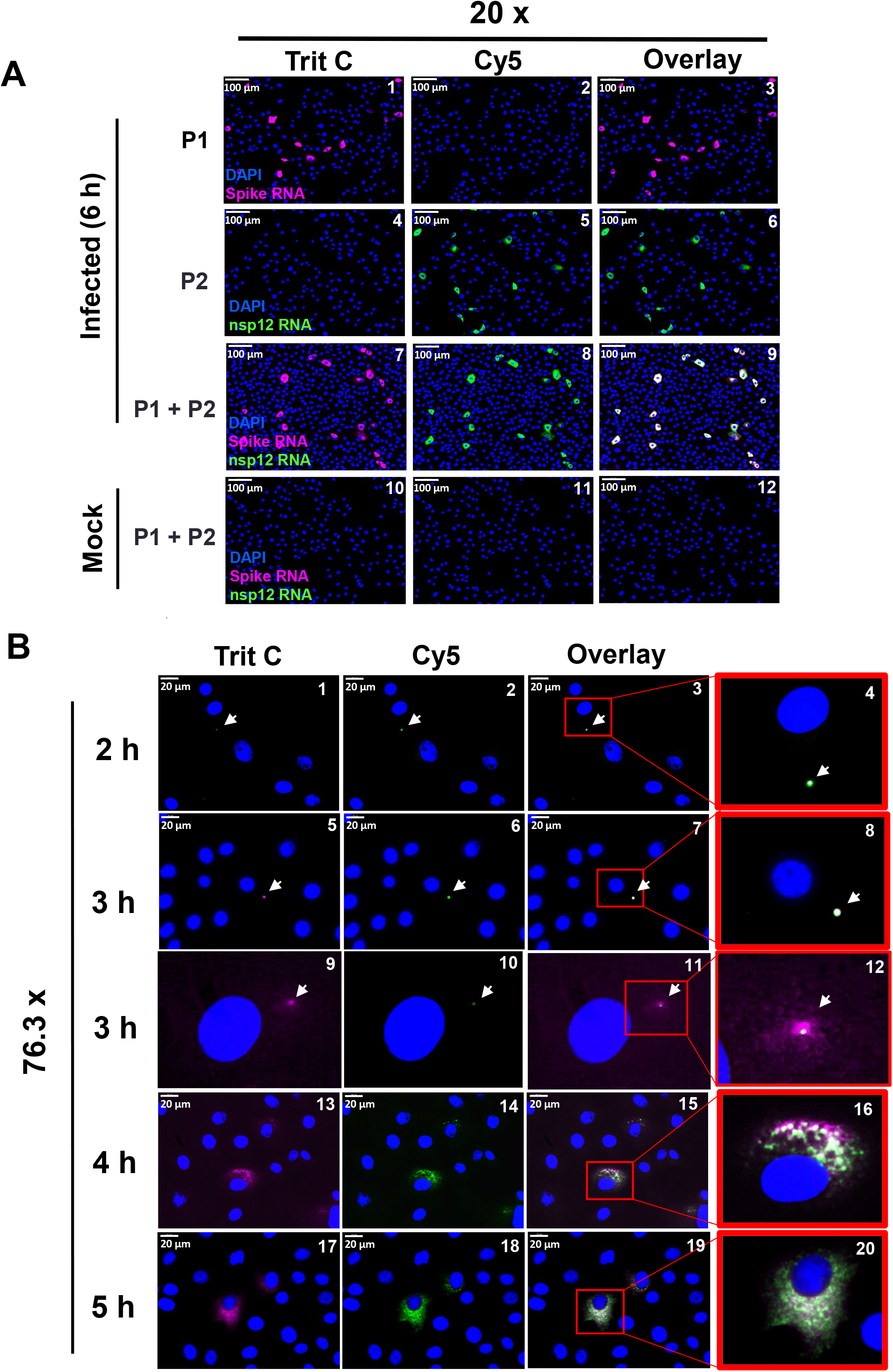
Simultaneous detection of SARS-CoV-2 gRNA and sgRNA-S in a time-course analysis. **(A)** Validation of spike RNA probe P1 and nsp12 RNA probe P2 to simultaneously detect gRNA and sgRNA-S in the infected cells. Cells were infected for 6 hours and hybridized with probes P1 and P2 alone or together and subjected to smRNA-FISH. Images are represented at 20x magnification. Scale bar at 100 μm. **(B)** A time-course analysis to detect the replication of gRNA and sgRNA in infected cells: The infected cells at 2, 3, 4 and 5 hours p.i. were hybridized with both spike RNA probe P1 and nsp12 RNA probe P2 and subjected to high-speed, high-resolution scanning. The images are represented at 76.3x magnifications. Panels 4, 8, 12,16 and 20 are the zoomed images of insets for indicated time-points. White arrows point to potential replication organelle (RO). Blue indicates DAPI stained nuclei. Scale bar at 20 μm.

We carried out another time course analysis using the two probes together to determine the early replication kinetics of gRNA and sgRNA-S. As before, we used a low m.o.i. of virus for infection with the purpose of following the replication of a single virus soon after its entry into the cell after 0.5, 1, 2, 3, 4, 5 and 6 hours p.i. Scanning of the entire slides containing the infected cells and analyzing the images at low magnification (5x), indicated the presence of strong positive cells at 4 hours p.i (S3 Fig; panels 16-18). The infected cells were positive for both gRNA and sgRNA-S (S3 Fig). As the time progressed, the number of cells that were positive increased at 5 and 6 hours p.i. (S3B Fig; panels 19-24). We did not observe positive cells within uninfected samples (S3 Fig; panels 25-27). Analysis of the above images at higher magnification (76.3x) indicated that a majority of the cells at 2 hours p.i., and many cells at 3 hours p.i. appeared to have a single RNA spot positive for both P1 and P2 probe, indicating that these spots represented viral gRNA at the earliest stages of its replication (Fig 3B; panels 1-8; white arrows). Because of resolution limitations using the scanner, the single positive RNA spots we identified are unlikely to be single RNA molecules, and it is likely that they represented groups of RNA molecules. At 3 hours p.i., the single RNA cluster/patch present within some cells appeared to contain both gRNA and sgRNA-S as distinct spots within the patch, and these spots exhibited variable degrees of hybridization to P1 and P2 probes (Fig 3B; panels 9-12). The spots at 3 hours p.i. were larger than the spots observed at 2 h p.i. (Fig 3B; panels 9-12 white arrow). At 3 hours p.i., within the single RNA clusters/spots, there was a central area positive for both P1 and P2 probes, which was surrounded by spots that hybridized to only P1 probe, indicating that perhaps the central region of these clusters likely harbor gRNA and the RNA molecules in the periphery of this central region corresponded to sgRNA-S (that hybridized to P1 probe only) that are migrating away from this central spot (Fig 3B; panels 9-12). This result suggested that these individual RNA clusters at early time points could represent virus induced RO that harbored genomic RNAs in the center surrounded by the sgRNA-S radiating and migrating out of these ROs. We surmise that the single spots per cell observed at 2 h and 3 hours p.i. likely represent the formation of a first RO by the SARS-CoV-2 RNA that entered the cell. At 4 and 5 hours p.i., the infected cells harbored multiple viral RNA spots that were visualized as large puncta of varied size and number (Fig 3B; panels 13-20).

To gain further insight into the SARS-CoV-2 positive spots that were observed at early time points (2-3 hours p.i.) in infected cells, the above samples were subjected to microscopy for detecting single molecules as described (Fig 4). We found that at 2 hours p.i. many cells harbored either one or two single RNA molecules that uniformly hybridized to both P1 and P2 probes, confirming that at this time point, only gRNAs but not sgRNAs were present (Fig 4; panels 5-8, White arrows). At 3 hours p.i. we found an increase in the number as well as size of spots that hybridized to P1 and P2 probes (Fig 4; panels 9-16). The size of the spots within the same cell varied considerably, indicating that these spots represented clusters of RNA at various stages of replication (Fig 4; Panels 12 and 16). As we had noticed in the low-resolution analysis, the central areas in these spots hybridized to both P1 and P2 probes, representing gRNA, which were surrounded by radiating and migrating spots that hybridized to only P1 probe, representing the sgRNA-S (Fig 4; panels 9-16). The size of the peripheral spots around the RO that hybridized to P1 probe alone were similar in size to that of a single RNA molecule found at 0.5 hours and 2 hours p.i. (Fig. 4; Panels 14-16, white arrows point to RO and yellow arrowhead points to single sgRNA-S molecules). We observed that the number and size of the positive spots increased at later time points and the intensity with which these spots hybridized to P1 and P2 probes varied (Fig 4; panels 17-28, and Fig 5; panels 1-8).

**Fig 4.**
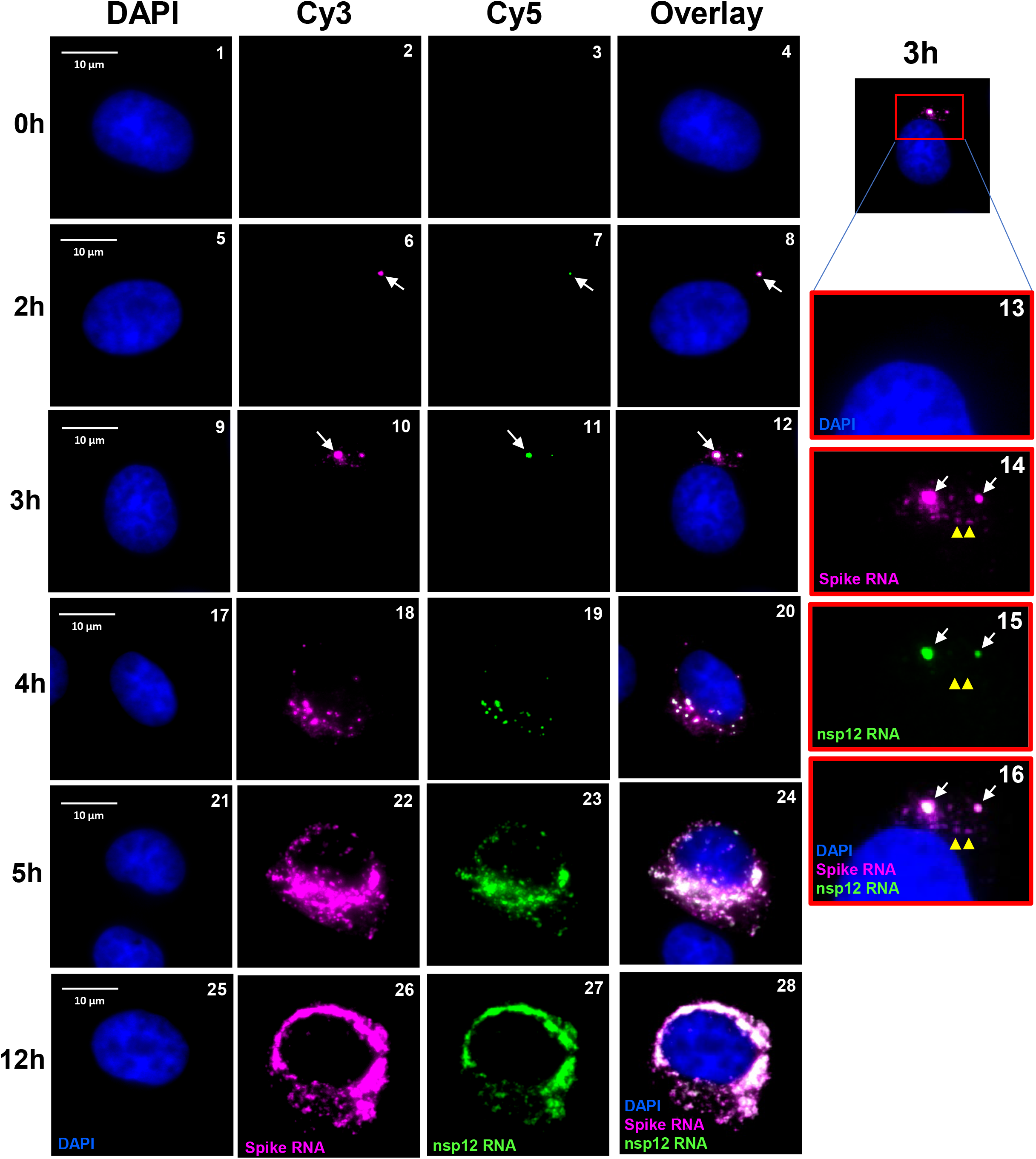
Single molecule analysis to determine the kinetics of SARS-CoV-2 gRNA and sgRNA replication. Transcript specific visualization of gRNA and sgRNA-S in SARS-CoV-2 infected Vero E6 cells using P1 (spike, represented in magenta) and P2 (nsp12, represented in green) probes. When spike RNA probe P1 and nsp12 RNA probe P2 both hybridize to the same molecule, it is shown in white (overlay). Blue indicates DAPI stained nuclei. Images represent the cells infected with SARS-CoV-2 and hybridized with P1 and P2 probes at 0, 2, 3, 4, 5 and 12 hours p.i. White arrows in panels 6-16 point to RO. Yellow arrow heads point to single molecules of sgRNA-S. Panels 13-16 show the zoom of the corresponding overlay images at 3 hours p.i., illustrating two ROs (white arrows). The larger RO in these panels harbors gRNA in the center surrounded by the sgRNA-S. Single molecule RNA are indicated by yellow arrow heads (panels 14-16). The images were acquired using a wide-field microscope. Scale bar at 10 μm.

**Fig 5.**
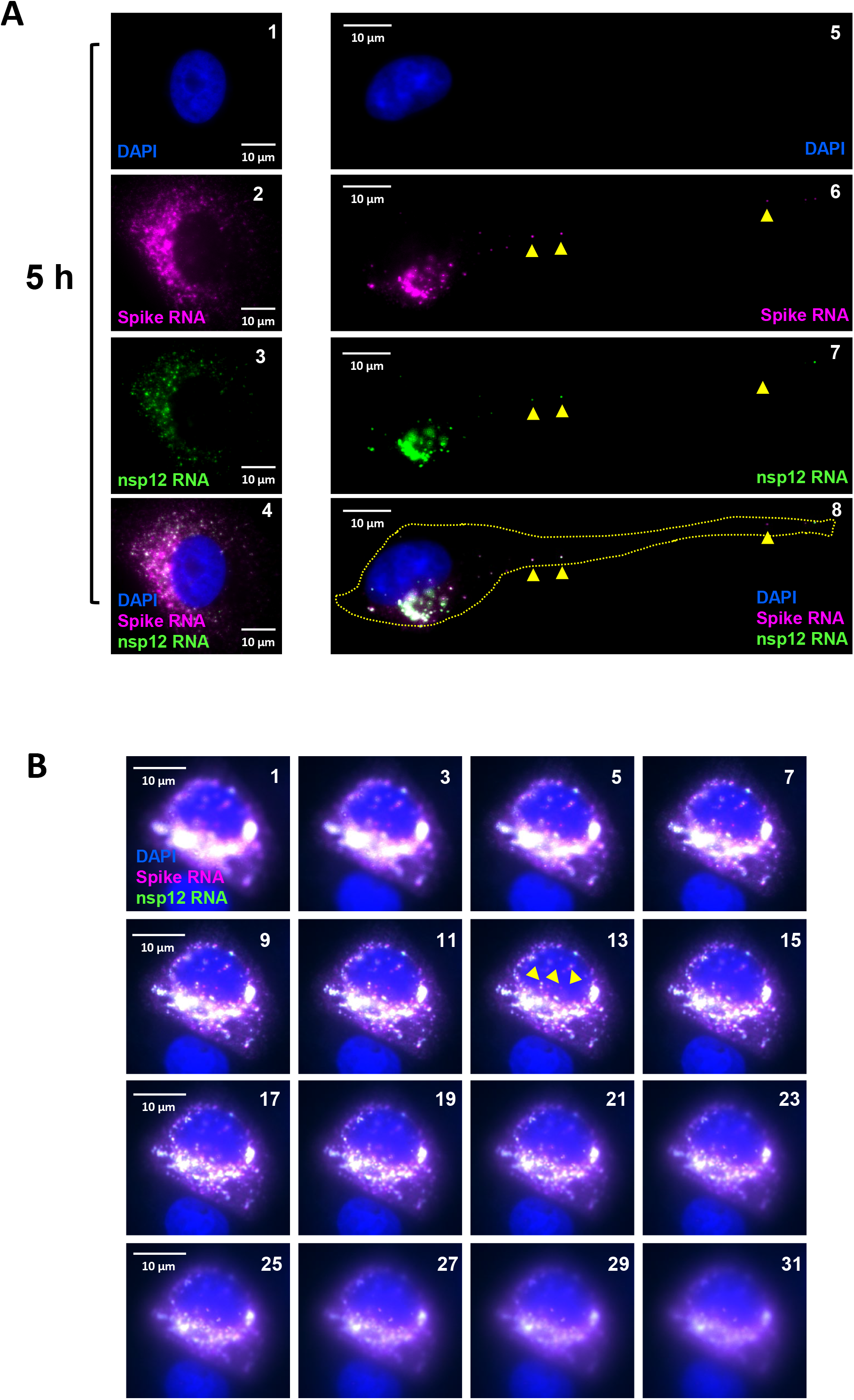
Images of cells infected with SARS-CoV-2 at 5 hours p.i. **(A)** Representative cells infected with SARS-CoV-2 RNA at 5 hours p.i. Panels 1-4 represent a cell filled with ROs at 5 hours p.i. Panels 5-8 represent a single cell with migrating gRNA and sgRNA-S to the distal end of the cell (yellow arrow heads). **(B)** Presence of SARS-CoV-2 RNA in the nucleus. The images from a few of the Z-stack of cell are represented to demonstrate the presence of the positive RNA spot inside the nucleus. The Z-stack images were acquired starting from the bottom of the cell moving upwards. The panel numbers represent the stack number in a total of 42 stacks. In both A and B, images were acquired using a wide-field microscope. The spike RNA P1 probe signals are colored in magenta (sgRNA-S), nsp12 RNA P2 probe signals are colored in green (gRNA) and when P1 and P2 both hybridize to the same molecule, it is shown in white (overlay). Scale bar, 10 μm. DAPI (nucleus, blue).

These results suggested that the individual positive spots at 2 hours p.i. and earlier time points likely represented single gRNA molecules within each cell, but the larger spots at 3 hours p.i. and at later time points represented RO with gRNA in the center surrounded by sgRNA-S in the periphery. The time-course experiments demonstrated that the single genomic RNA that entered the cell is preparing to replicate for about 1-2 hours p.i. By ~3 hours p.i. the formation of RO from single gRNA allows rapid replication of gRNA and transcription of sgRNAs. After 3 hours p.i. the formation of a single parental RO is followed by the formation of other daughter ROs, as indicated by the increase in number and size of RO per cell (Fig 4, panel 12 and panels 14-16; white arrows). In some cells we noted that both gRNA and sgRNA were migrating away from the nuclear periphery and reached the narrow and elongated distal regions of the cells (Fig 5A; panels 5-8; yellow arrow heads). At later time points of 5-12 hours p.i., the ROs spread-out from nuclear periphery, filling the entire cytoplasm (Fig 4; panels 21-28, and Fig 5A; panels 1-4). Interestingly, we also observed SARS-CoV-2 positive spots in the nucleus, the significance of which is unclear at this point (Fig 4; panels 21-24 and Fig 5B, yellow arrow heads).

### Heterogeneity in the replication of SARS-CoV-2 RNA

Our results above indicated that the ROs can be distinguished by clusters of RNA by the presence of central gRNA surrounded by the sgRNA-S. Furthermore, upon examination of large number of infected cells scanned using HSHRS-FM, we noted that there is a heterogeneity in the appearance of RNA in the infected cells at any given time point (S4 Fig). At early time points majority of the infected cells had either one or a few ROs per cell and at later time points majority of infected cells had large number of ROs filling the entire cytoplasm (S4 Fig). To quantitate the progression of replication of RNA through different stages, we used images obtained with P2 probe, as this probe hybridizes to genomic RNA and not to sgRNA making it easier to identify individual RO as speckles. We arbitrarily defined various stages of replication as follows. Stage-I replication was defined as when cells harbored 1-5 speckles of RO per cell but not much diffused RNA (Fig 6A, panels 1 and 2). Stage-2 and Stage-3 were defined as intermediates where number of ROs progressively increased (Fig 6A, panels 3-6). Stage-4 was defined as a late stage when the entire cytoplasm was filled by ROs (Fig 6A panels 7 and 8). We analyzed ~1000 cells in 10 different random fields of views were acquired using the HSHRS-FM per time point and the positively infected cells containing virus at various stages of replication were quantitated at 4, 5 and 6 hours p.i. and plotted as fraction of positive cells at various stages at a given time point (Fig 6B). The results indicated that at 4 hours p.i., approximately 35% of the infected cells showed viral replication at Stages 1-3 and a few cells (2.5%) at Stage-4 were seen. At 5 hours p.i. the Stages 1-3 decreased to 14 to 26% and Stage-4 increased to 43%. There were very few cells with Stage-4 (2.5%) at 4 hours p.i., which gradually increased (51%), culminating at 6 hours p.i. These data clearly indicate that the virus replication progresses from a single RO to multiple ROs per cell and that it is asynchronous or heterogeneous.

**Fig 6.**
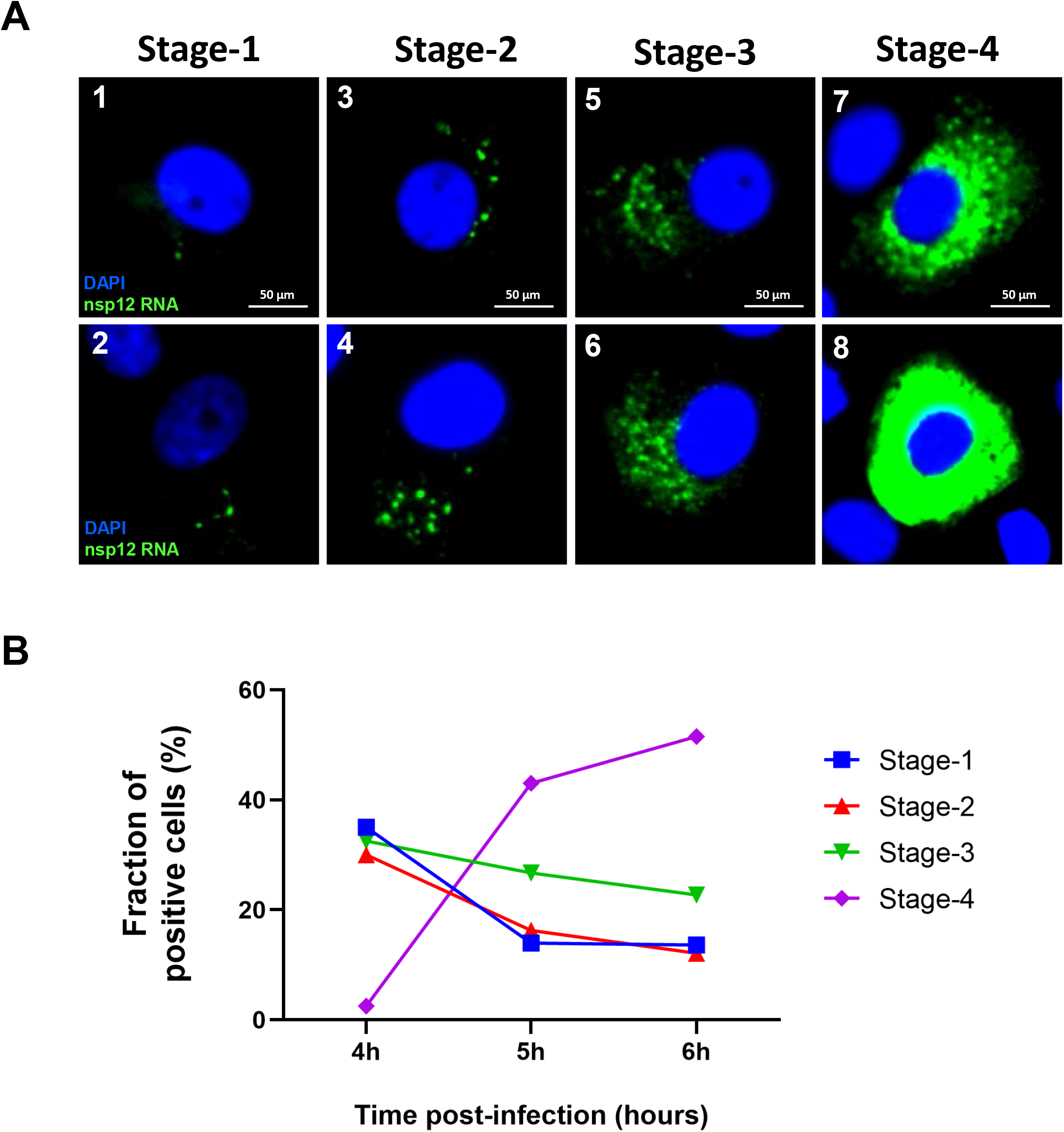
Quantitation of cells containing virus at different stages of replication. **(A)** Examples of images of cells with four different stages of viral replication at Stage-1 (panels 1 and 2), Stage-2 (panels 3 and 4), Stage-3 (panels 5 and 6), and Stage-4 (panels 7 and 8). Blue color represents DAPI nuclear staining; Green color represents ROs containing genomic RNA. Scale bar, 50 μm. **(B)** Graphical representation of quantitation of percentage of positive cells harboring four different stages of replication at 4, 5 and 6 hours p.i.

## Discussion

Our studies provide a spatial and temporal characterization of early post entry events of SARS-CoV-2 RNA replication in Vero E6 cells at a single cell, single molecule resolution, and sheds light on the formation and progression of an RO originating from a single RNA molecule that entered the cell. Our report provides information about localization of gRNA and sgRNA within an RO and demonstrates the intracellular dissemination of the sgRNA-S from an RO within the cytoplasm of an infected cell. Our data also indicate that after the formation of the first RO, additional ROs are formed within a short period of time, which further increase in number to entirely fill the infected cell cytoplasm. We also noted that there is a cell-to-cell heterogeneity in the rate of replication of RNA as can be observed by quantitating the number and nature of ROs present within cells at any given time point. Different % of cells containing RNA at different stages of replication can be found at all time points while at early time points most of the cells have RNA at the early stages of replication, at later time points, most of the cells have RNA at a later stage of replication (Fig 6). As the time progresses, percentages of cells with early stages of RNA replication decreases and percentages of cells with late stages of replication increases. This indicates that there was a randomness in which the rate of replication was progressing. The reason for different rates of replication in different cells is not clear. The heterogeneity in replication could be due to differences in entry of RNA into the cytoplasm, establishment of the first RO from the first RNA that entered cell, or it could be due to differences in the rate at which the first RO gives raise to multiple ROs. It is well known that stage of cell cycle and other cellular factors affect the rate of viral replication [21, 22]. Based on our observation, the heterogeneity appears to be a random or stochastic event, as all stages of replication are found at any given time point.

We detected cytoplasmic localized single gRNA molecules per cell at 0.5 hours, 1 hours and 2 hours p.i. and these molecules likely represent viral RNA of a particle that has just entered the cell. To our knowledge, these represent the earliest time points of detection of SARS-CoV-2 viral RNA in an infected cell. Our studies also indicated that a single RNA molecule forms distinct viral replication center that eventually appear as clusters of RNA, likely representing an RO. These RO begin to appear as early as 2 hours p.i. and by 3 hours p.i., the ROs harbored multiple gRNAs in the center surrounded by sgRNA transcripts that radiate and migrate out of these centers. We surmise that as the replication of gRNA continues within the first RO, some of these gRNAs migrate out to form “daughter” ROs in the same cell. Thus, we predict that kinetics of replication of the positive strand RNA into negative strands and replication of these negative strands into additional gRNA and sgRNAs can be initiated between 0.5 hours and 2 hours p.i. At later time points, multiple ROs are formed within the same cells and by 5 hours p.i. most of the cytoplasm is filled with ROs loaded with SARS-CoV-2 RNA and by 6 hours, it is hard to distinguish individual ROs within the same cell. These results establish the time-course of early replication, allows the earliest visualization (0.5 hours and 1 hours p.i.) of SARS-CoV-2 replication inside the cells and indicate that a single gRNA begins to form a distinct RO at ~2 hours p.i. Incidentally, we detected the presence of molecules of SARS-CoV-2 gRNA within the nuclei of some of the cells. However, at this point, we do not know if these nuclear viral RNA are significant for replication. It is possible that these RNA are not in the nucleus but are in the cytoplasmic invaginations that are separated by nuclear membrane. More experiments are needed to establish the existence of nuclear speckles of SARS-CoV-2 RNA. In summary, high resolution early kinetic analysis of SARS-CoV-2 RNA replication provides an understanding of the timing of formation and arrangement of gRNA and sgRNA within an RO and sets the stage for further analysis of these unique organelles in the future.

While our manuscript was in preparation, Lee et. al. reported the use of smFISH to study the infection of SARS-CoV-2 in human cell lines [14]. They simultaneously visualized and quantified positive-sense along with negative-sense viral RNA genomes and sub-genomic RNAs. They were able to detect the viral infection 2 hours p.i. and observed the initial production of sub-genomic RNA within the RO. While our data is consistent with the above report, we were able to identify viral RNA within the cells at earlier time points (0.5 hours and 1 hours p.i.) and were able to image ROs or viral RNA factories at much higher resolution, where we identified the presence of a central region within these ROs filled with gRNA that are surrounded by migrating sgRNAs. Lee et al. also quantitated individual RNAs within replicating cells. The fact that ROs begin to form as early as ~2 hours p.i. suggests that it is hard to find individual gRNAs after 2 hours p.i. We believe that the presence of numerous gRNAs and sgRNAs within each individual ROs makes it difficult to get precise quantitation of RNA molecules per cell. Therefore, it is likely that absolute quantitation of single RNA molecules after 2 hours p.i. reported by previous studies, is underestimated [14].

Our study establishes the use of RNA-FISH technology in assessing the early kinetics of SARS-CoV-2 replication. These studies will provide a platform for addressing several emerging questions related to mechanism of SARS-CoV-2 replication. Furthermore, the assays we have developed could be used in the future to study the formation of replication organelles, involvement of viral proteins in ROs and the kinetics of formation of RO in relation to the timing of synthesis of replication intermediates. Such studies could help identify new and unique therapeutic targets to combat SARS-CoV-2 replication.

We are currently optimizing methods to detect single molecules in tissues from autopsies, lavages or sputum from COVID-19 patients to facilitate the detection of low levels of SARS-CoV-2 replication in a tissue- and organ-specific manner, which is beyond the scope of this manuscript. We believe that the methodologies that we develop may address the long-standing questions about post-COVID and long-COVID. Furthermore, the technology reported here could be used in patient samples to determine responses to therapeutic drugs.

## Materials and Methods

### Cell culture

Vero E6 cells (ATCC, CRL-1586) were maintained in Dulbecco’s Modified Eagle Medium (Cat: SH30081.01; Hyclone) supplemented with 10% fetal bovine serum, 2mM L-glutamine, non-essential amino acids, 100 U/mL penicillin and 100 μg/mL streptomycin. All the cell lines were maintained in a standard 5% CO_2_ in a culture incubator at 37° C.

### Infection of Vero E6 cells with SARS-CoV-2

For infection of Vero E6 cells with SARS-CoV-2 (SARS-CoV-2 USA-WA1/2020), cells were seeded on 4-well chamber slides (Nunc) at a confluency of 50-70% the day before infection. On the day of the infection, medium was replaced with 500 μl of infection medium (DMEM medium supplemented with 2% fetal bovine serum, non-essential amino acids, HEPES and penicillin/streptomycin) and the cells were transferred to the BSL-3 laboratory. The virus was diluted in infection medium and 100 μl/well added to each well to achieve a m.o.i. of 0.5 PFU/cell. After, 0, 0.5, 1, 2, 3, 4, 5, 6,12 and/or 24 hours post-infection, supernatant was removed and the cells were washed twice with 1 ml of 1x PBS with 5 mM MgCl_2_. After the second wash, cells were fixed with 4% paraformaldehyde. During fixation, cells were protected from light.

### Immunofluorescence

The fixed cells were washed with 1x PBS, permeabilized with 0.2% Triton X-100 and incubated with a mouse monoclonal antibody that recognizes the NP protein of SARS-CoV-2 (1C7, kindly provided by Dr. Thomas Moran, Icahn School of Medicine at Mount Sinai). Next, cells were washed with 1x PBS and incubated with Alexa fluor 488 conjugated anti-mouse secondary antibody (Invitrogen) and DAPI. The signals were detected using an EVOS M5000 fluorescent microscope.

### smRNA-FISH probe design and specificity analysis

Forty different 22 nucleotide long smRNA-FISH probes (5’-> 3’) for spike (S) and RdRp (nsp12) genes were generated using LGC Biosearch Technologies’ Stellaris® RNA FISH Probe Designer version 4.2 [23]. Each probe for spike gene was tagged with Quasar 570 and for nsp12 gene with Quasar 670 dyes at 3’ ends, respectively. As a target reference sequence, coding sequences (CDS) of Spike and nsp12 region was selected from the SARS-CoV-2 Wuhan-Hu-1 (NC_045512.2) reference sequence. Each probe sequence was subjected to BLAST and was screened against other coronavirus sequences, human transcriptome, and human intron database. To perform in silico probe sequence specificity analysis, all 40 oligonucleotide sequences of the Spike and nsp12 smRNA-FISH probes were aligned against; SARS-CoV-2 (NC_045512.2); SARS-CoV-1 Tor2 (NC_004718.3); MERS-CoV isolate HCoV-EMC/2012 (NC_019843.3); HCoV-HKU1 (NC_006577.2); HCoV-OC43 strain ATCC VR-759 (AY585228.1); HCoV 229E strain 229E/human/USA/933-40/1993 (KF514433.1); HCoV NL63 strain NL63/human/USA/0111-25/2001 (KF530112.1) and Human hg38_mRNA (AF001540.1), RefSeq genome or transcriptome assembly, using ‘bowtie2’ (2.4.4). To get the minimum edit distance of oligonucleotide sequences to target genome/transcriptome, following bowtie2 v2.4.4 arguments were used; --end-to-end, --no-unal, --align-seed-mm 0, --align-seed-length 5, --align-seed-interval 1-1.15, -- effort-extend 15, --effort-repeat 2 [14]. The heatmaps were created using R v4.0.2 with Bioconductor package Complex Heatmap v2.9.3 [24].

### Single molecule RNA fluorescence in situ hybridization (smRNA-FISH)

Vero E6 cells (ATCC, CRL-1586) seeded on a four-chambered slide were inoculated with SARS-CoV-2 (USA-WA1/2020) at an m.o.i. = 0.5, for various time points (0, 0.5, 1, 2, 3, 4, 5, 6,12 and/or 24 hours). The cells were fixed with 4% paraformaldehyde post-infection for 30 minutes followed by smRNA-FISH analysis. After fixation, each well of the four chambered slide was treated with 0.1 M Glycine/PBSM for 10 min at room temperature followed by cell permeabilization with PBSM/0.1% Triton X-100 for 10 min. After washing with 1x PBSM, cells were pre-hybridized in 2x SSC, 15% formamide for 30 min at 37° C. The cells were incubated overnight with 300 μl hybridization buffer containing SARS-CoV-2 spike and/or RdRp/nsp12 stellaris probes tagged with Quasar 570 and Quasar 670 dyes, respectively. 25 μM of SARS-CoV-2 spike and RdRp/nsp12 stellaris probes (1 μl probe/100 μl hybridization solution) were added to hybridization buffer containing, 10% dextran sulfate (Cat: D61001; Sigma), 1mg/mL tRNA from *E. coli* MRE 600 (Cat No: 10109541001; Millipore Sigma), 15% formamide (Cat: 75-12-7; Acros Organics), 0.2 mg/mL UltraPure BSA (Cat: AM2616; Ambion™), 2x Sodium Chloride-Sodium Citrate (SSC) buffer (Cat: R019, G-Biosciences), 2 mM Vanadyl Ribonucleoside Complex (Cat: S1402S; New England Biolabs), and 10 U/mL SUPERase•In RNase Inhibitor (Cat: AM2694; Ambion™). The slides were incubated in a humid hybridization chamber overnight at 37° C. After the overnight hybridization, cells were washed with pre-hybridization buffer (2x SSC, 15% formamide) three times followed by 3-4 times washing with 2x SSC. The cells were stained with DAPI (1 μg/ml in 2x SSC) for 2 min at RT. After DAPI staining the cells were washed with 2x SSC and mounted using the ProLong Gold antifade mountant (Cat: P10144, Thermo Scientific).

### Whole-slide scanning, Microscope setup and image acquisition

Fluorescent scanning of the slides was done in two stages. First, whole-slide scanning of the entire slide was obtained at 20x using PANNORAMIC 250 Flash III Slide Scanner. Automatic focusing (default factory settings) was used for scanning the whole slide. For fluorescence imaging, we used DAPI, Cy5-Q and TRITC-Dendra fluorescence filters. Pannoramic scanner software were used for image acquisition and scanned slides were visualized using CaseViewer 2.4 (64-bit version). For setting the parameters for image acquisition, mock control was used to set the background level of the fluorescence in the test images. After scanning, the same slides were subjected to single cell imaging. Cells were imaged using an upright, wide-field Olympus BX-63 Microscope equipped with a SuperApochromatic 60×/1.35 NA Olympus Objective (UPLSAPO60XO), a SOLA light engine (Lumencor), an ORCA-R2 Digital Interline CCD Camera (C10600-10B; Hamamatsu), and zero-pixel shift filter sets: DAPI-5060C-Zero, Cy3-4040C-Zero (for Quasar 570 detection), and Cy5-4040C-Zero (for Quasar 670 detection) from Semrock, as described [25]. The resulting image pixel dimension was 107.5 nm, and the z-step size (along the optical axis) used for all optical sectioning acquisition was 200 nm. Metamorph software (Molecular Devices) was used for controlling microscope automation and image acquisition. Images were analyzed using ImageJ and/or Fiji software [26].

## Author contributions

**Conceptualization**: Ganjam Kalpana.

**Funding acquisition**: Ganjam Kalpana, Adolfo García-Sastre, Carolina Eliscovich

**Investigation**: Rajiv Pathak, Carolina Eliscovich, Ignacio Mena, Updesh Dixit, Adolfo García-Sastre, Robert H Singer, Ganjam V. Kalpana.

**Methodology**: Rajiv Pathak, Carolina Eliscovich, Ignacio Mena, Updesh Dixit.

**Supervision**: Ganjam Kalpana, Robert Singer, and Adolfo García-Sastre.

**Visualization**: Rajiv Pathak and Carolina Eliscovich.

**Writing-original draft**: Ganjam Kalpana and Rajiv Pathak with contribution from other authors.

**Writing-review and editing**: Rajiv Pathak, Carolina Eliscovich, Ignacio Mena, Adolfo García-Sastre, Robert Singer, Ganjam Kalpana.

## Acknowledgments

We thank Dr. V. Prasad for critically reading the manuscript. We thank Drs. Kartik Chandran and Rohit Jangra for kind gift of VSV-virus containing codon optimized Spike protein ORF. The current work was in part funded by NIH grant 1 R01 DA 043169-01 to GVK as MPI; NIH grant R01 DK110063 to CE; and CRIPT (Center for Research on Influenza Pathogenesis and Transmission), a NIAID funded Center of Excellence for Influenza Research and Response (CEIRR, contract # 75N93021C00014) to AG-S. We thank Andrea Briceno at Albert Einstein College of Medicine Analytical Imaging Facility (AIF), for assistance with scanning of images on PANNORAMIC 250 Flash III Slide Scanner. AIF was supported by NIH grant 1S10OD026852-01A1 and Cancer Center Grant P30CA013330. We also thank Randy Albrecht for support with the BSL3 facility and procedures at the Icahn School of Medicine at Mount Sinai (ISMMS) and Richard Cadagan for excellent technical assistance.

## Supporting information

**S1 Fig. Specificity of SARS-CoV-2 spike RNA probe P1 in detecting SARS-CoV-2 viral genome. (A)** Vero E6 cells mock-infected or infected with VSV-Spike containing codon-optimized spike gene open reading frame, were probed with spike RNA probe P1 after 24 hours p.i. The cells were imaged under FITC channel for expression of GFP (shown in green, panels 1 and 3) or under TritC channel for detecting hybridization with P1 probe (magenta, panels 2 and 4). GFP expression demonstrates the infection of cells (green, panels 1 and 3), whereas the absence of a P1 probe signal (magenta) with cells infected with VSV-Spike, indicates the Specificity of SARS-CoV-2 spike RNA probe P1. **(B)** Vero E6 cells mock- or SARS-CoV-2 (USA-WA1/2020)-infected, at 24 hours p.i. (m.o.i.: 0.5 PFU/cell), were probed with spike RNA probe P1 (magenta, panels 1 and 2). The blue color indicates nuclei upon DAPI staining, magenta indicates hybridization with P1 probe. The scale bar is 150 μm.

**S2 Fig**. **Presence of SARS-CoV-2 RNA within the cytoplasm and nuclei of infected cells 30 min p.i. SARS-CoV-2 RNA in the cytoplasm**. **(A)** and in the nucleus. **(B)** The panels represent Z-stack images of an infected cell to demonstrate the presence of the positive RNA spot inside the cell. A total of 42 Z-stacks were acquired using widefield microscopy starting from the bottom of a cell moving upwards. The panel numbers in the figure represent the stack number of the Z-stacks. Blue color represents DAPI staining and Magenta represents SARS-CoV-2 spike RNA. The scale bar is 10 μm.

**S3 Fig. A time course analysis of SARS-CoV-2 replication to simultaneously detect gRNA and sgRNA-S**. Vero E6 cells were infected with SARS-CoV-2 and hybridized with probes at 0, 0.5, 1, 2, 3, 4, 5 and 6 hours p.i. The infected cells were probed using both spike RNA probe P1 and nsp12 RNA probe P2. Four chambered slides containing infected cells and uninfected controls, probed with spike RNA probe P1 and nsp12 RNA probe P2, were subjected to high-speed, high-resolution scanning. The panels represent images of an entire well of the infected cells at 5x magnifications. Blue represents DAPI staining, green represents gRNA detected by nsp12 RNA probe P2, and magenta represents sgRNA-S detected by spike RNA probe P1and the overlay of the two probes shown in white. The scale bar is 500 μm.

**S4 Fig. A representative image of SARS-CoV-2-infected Vero E6 cells demonstrating various stages of viral replication**. Vero E6 cells infected with SARS-CoV-2 were subjected to RNA-FISH at 6 hours p.i. using nsp12 RNA probe P2 that detects gRNA. The images were acquired using HSHRS-FM. The illustration is a representative image of a field view of cells at 40x magnification, showing different stages of the viral replication. The numbers in the image refer to the stages of replication, stages 1-4, respectively. Blue color refers to DAPI staining to indicate the nucleus, and the green color represents gRNA hybridized with P2 probe. The scale bar is 20 μm.

