## Supplemental Figures for "Visualization of early RNA replication kinetics of SARS-CoV-2 by using single molecule RNA-FISH"

#### Supplementary Figures

### Supplementary Figure S1

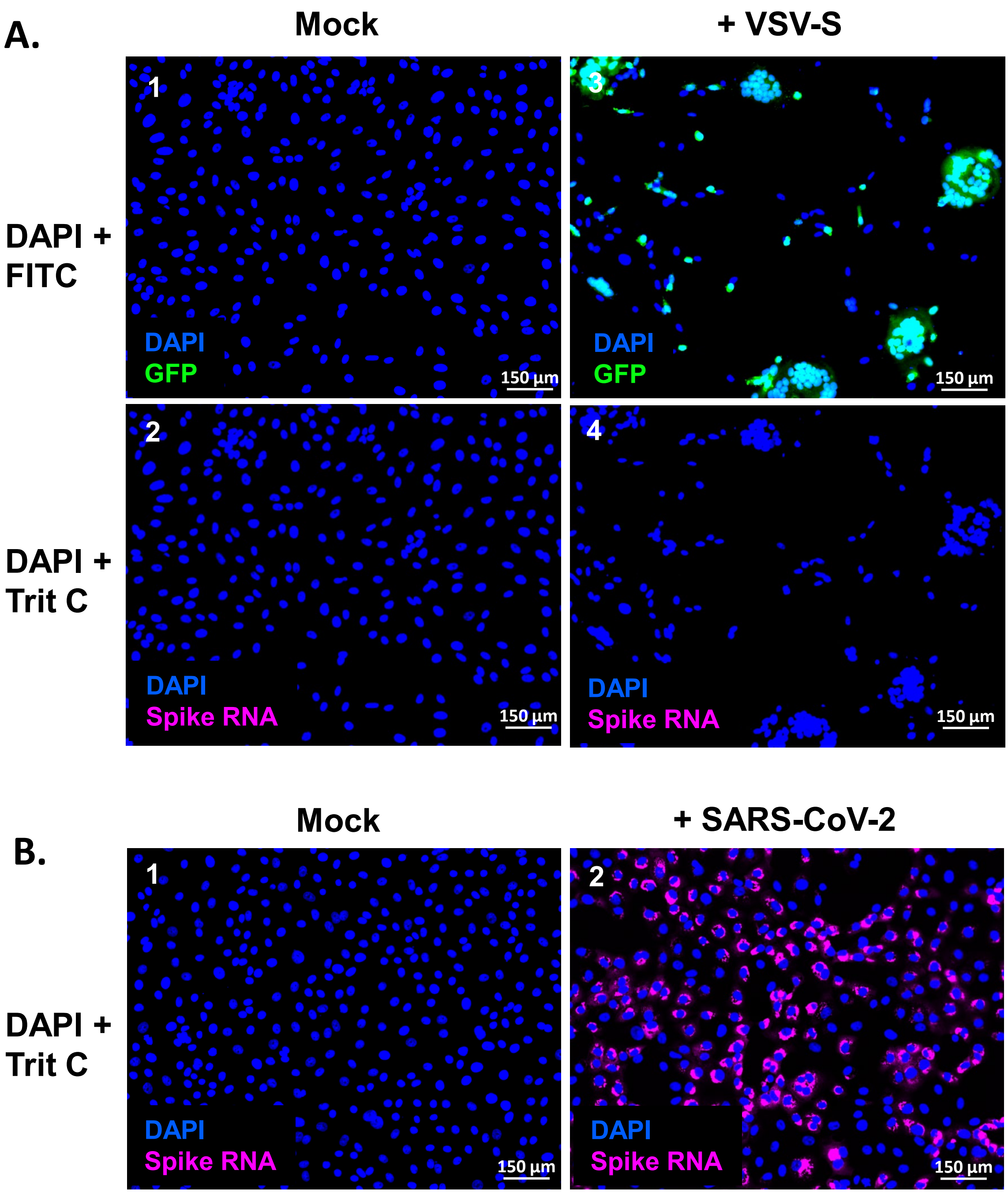

### Supplementary Figure S2

A.

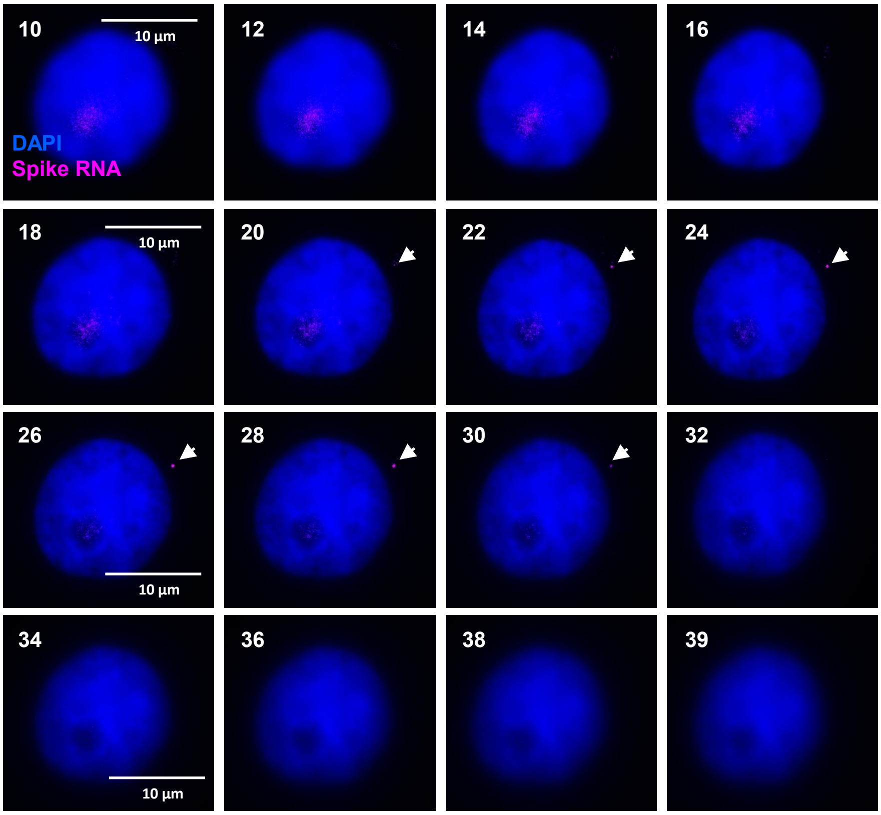


B.

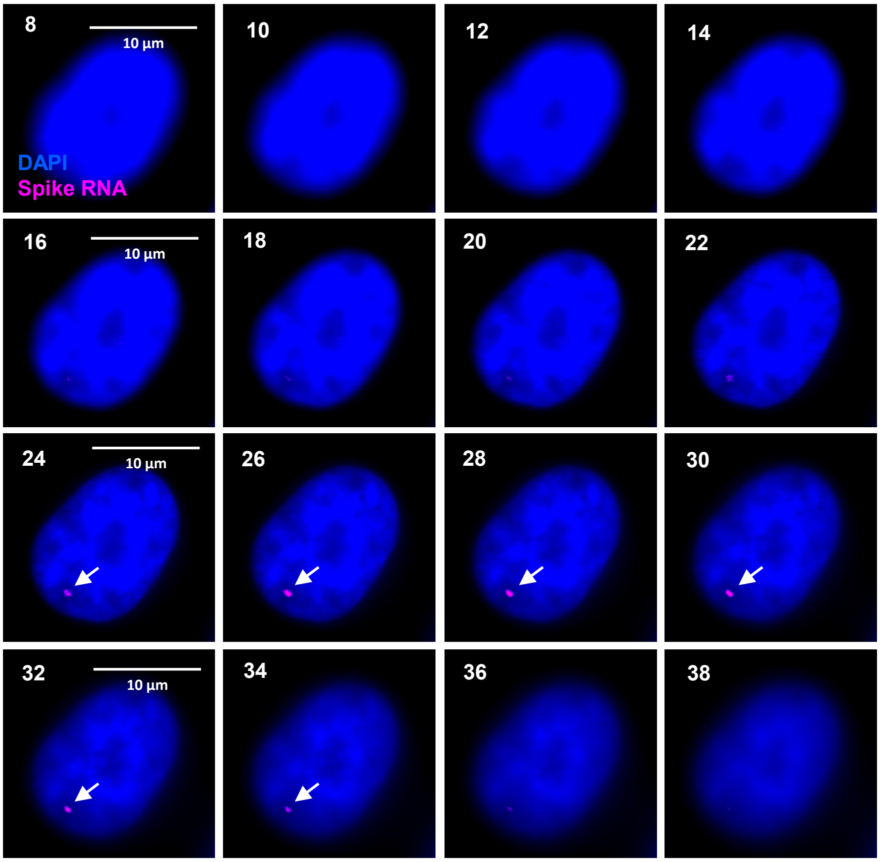

### Supplementary Figure S3

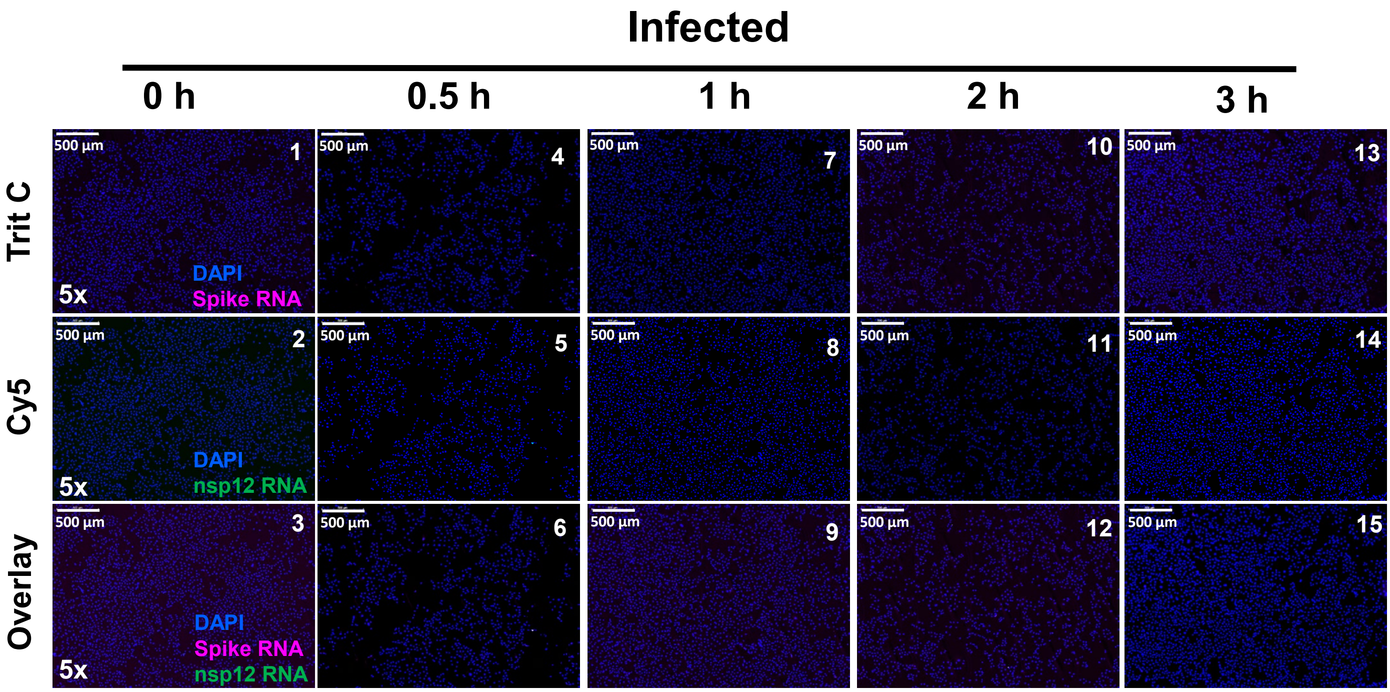


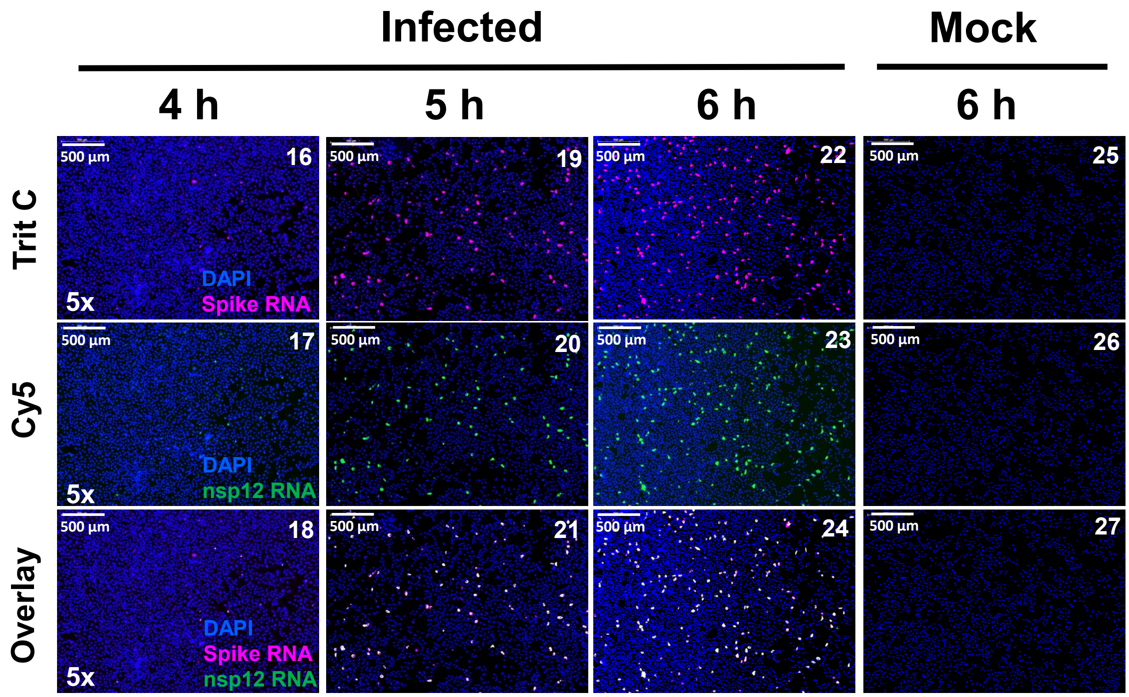

### Supplementary Figure S4

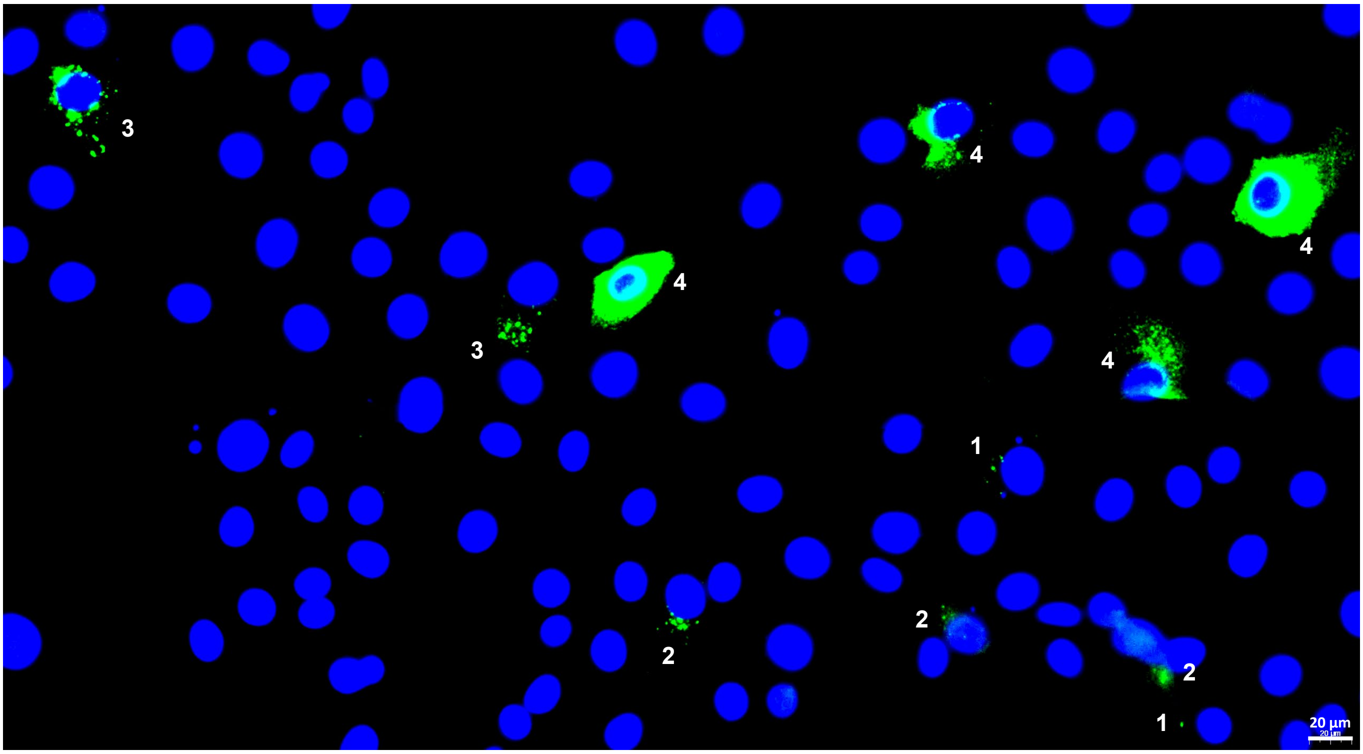
